# Cell-type-specific aberrant R-loop accumulation regulates target gene and confers cell-specificity

**DOI:** 10.1101/2022.07.19.500727

**Authors:** Xingxin Pan, L. Frank Huang

## Abstract

Aberrant R-loops have been found associated with diverse biological dysfunction, including cancers and neurological disorders. However, there isn’t any systematic research to characterize aberrant R-loops at the whole genome level at a large scale. Here, we identified aberrant R-loops, including proliferative and suppressive R-loops of 5’ end, body, 3’ end respectively for the first time, which are found prevalent and vary across diverse physiological conditions. After that, we proposed a deep neural network-based framework, named Deep R-looper Discriminant to identify aberrant R-loops against housekeeping R-loops. To evaluate the predictive performance of the deep learning framework, we constructed multiple prediction models as benchmarks and it showed our framework achieves robust performance for identifying aberrant R-loops against those normal R-loops. Furthermore, we found the customized Deep R-looper Discriminant was capable of distinguishing between proliferative and suppressive R-loops at 5’ end, body, 3’ end respectively, outperforming baselines. When inspecting the contribution of epigenetic marks to aberrant R-loops of each class, we inferred landmark epigenetic modifications which play a crucial role in the differentiated formation of those aberrant R-loops, and cell line specificity of epigenetic marks map was found as well. To explore the characteristics of these aberrant R-loops, we depicted the histone landscapes for aberrant R-loops. Finally, we integrated omics and identified target genes regulated directly by aberrant R-loops and found key transcription factors involved in R-loop regulation, which may be implicated in conferring cell-specificity and cancer development and progression.

## Introduction

R-loop is one kind of three-stranded structure that forms during transcription when the nascent RNA reanneals to the template DNA strand and leaves the non-template DNA strand alone[1]. Although R-loops occur over tens of thousands of genomic loci, there are mainly two R-loop hotspots: The promoter region is prone to R-loop formation where R-loops tend to peak 1-2kb downstream of the transcription start site (TSS) within the promoter-proximal transcription complex pausing sites; Terminal region is another R-loop hotspot where R-loop is prone to form surrounding transcription termination site (TTS)[2-5].

R-loop is significantly associated with genomic instability and dysregulation of R-loop metabolism has been implicated in multiple disorders, particularly neurodegenerative syndromes and cancer development[6-12]. However, the genomic R-loop profiles under pathological conditions have never been researched, and how to define, identify, and characterize these aberrant R-loops remains a question in the community. Topoisomerase factors have been reported to prevent R-loop formation in the specific condition and Ribonucleases H and RNA/DNA helicases are found involved in mediating R-loop resolution[13-17]. Besides, estrogen signaling in breast cells was shown to result in a burst of R-loop formation at thousands of loci and replication-dependent DSB, indicating upregulation of transcription may lead to elevated genic R-loops loads[18]. However, R-loop profiling of patients with Aicardi-Goutières syndrome harboring RNASEH2 loss of function didn’t reveal significant changes in genic R-loops, implying the regulation and function of aberrant R-loops are complicated and condition-specific[19].

With technologies advancing, several high-throughput R-loop profiling methods in genomic DNA such as DNA: RNA immunoprecipitation coupled to high-through DNA sequencing (DRIP-seq) are proposed, providing us with available public R-loop profiling data under diverse conditions in human[3, 20-35]. Meanwhile, with the development of artificial intelligence and high-performance computing, machine learning has been applied successfully to computational biology, particularly genome and omics[12, 36-38]. For example, deep learning has been applied to identify genetic variants affecting gene expression, predict DNA promoter methylation, histone modifications in DNA sequence, and even drug response effectively, providing us with exceptional potential solutions in the genomics fields[39-42].

In this study, we collected public available R-loop profiling data and identified the aberrant R-loops, including proliferative and suppressive R-loops at 5’ end, body, 3’ end of transcripts across conditions. After that, we proposed a robust deep learning framework named Deep R-looper Discriminant and it proved to achieve excellent prediction performance in detecting aberrant R-loops against normal R-loops compared with multiple baselines such as k-nearest neighbors (KNN), linear discriminant analysis (LDA), logistic regression (LR), naive bayes (NB), and random forests (RF). Next, we further proved the Deep R-looper Discriminant was able to distinguish robustly between proliferative and suppressive R-loops at 5’ end, body, 3’ end respectively after training. Furthermore, when accessing the importance scores of multi-omics spatial features derived from our framework, we inferred landmark epigenetic modifications across genome locations which may contribute to the differentiated formation of proliferative and suppressive R-loops in the corresponding region. To explore the characteristics of these six classes of aberrant R-loops, we depicted the epigenetic landscape for these aberrant R-loops around R-loop hotspots, including TSS and TTS. Finally, we integrated omics data and identified target genes directly regulated by these aberrant R-loops and common transcription factors medicated by the R-loops. It was found those target genes of aberrant R-loops are involved in conferring cell specificity and even cancer development.

## Results

### Aberrant R-loops are prevalent and show cell specificity across cell lines

To identify aberrant R-loops across the whole genome systematically, we first defined three regions for each transcript, including 5’ end region ranging from upstream 300bp of TSS to downstream 500bp of TSS, body region ranging from downstream 500bp of TSS to upstream 300bp of TTS, and 3’ end region ranging from upstream 300bp of TTS to downstream 3000bp of TTS. After mapping the reads to each region, we normalized the read densities of each region as R-loop level covering the corresponding regions (Figure 1A).

**Figure 1.**
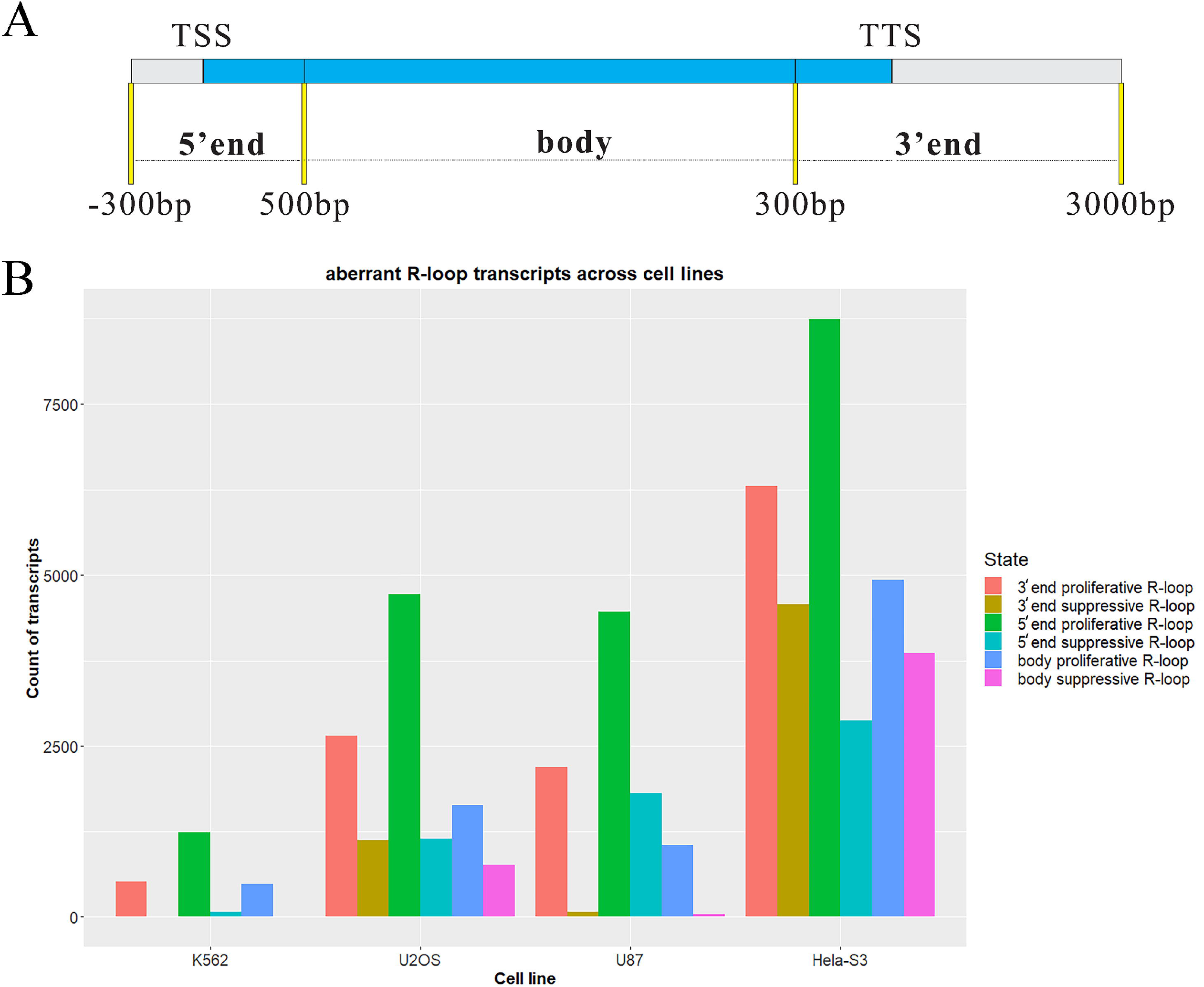
Aberrant R-loops identification. (A) The definition of aberrant 5’ end, body, 3’ end R-loops. (B) Aberrant R-loops identification across cell lines.

After obtaining R-loop level of all cancer cell lines, including K562, U2OS, U87, and Hela, we carried out differential R-loop analysis for these cancer samples against non-cancer samples including NT2, HEK293, and IMR90 via differential analysis based on the negative binomial distribution model of edgeR[43]. Thereafter, we identified aberrant R-loop transcripts for the cancer cell lines, including 5’ end proliferative R-loop, 5’ end suppressive R-loop, body proliferative R-loop, body suppressive R-loop, 3’ end proliferative R-loop, 3’ end suppressive R-loop, and normal R-loop (Figure 1B). Interestingly, proliferative R-loops predominate in aberrant R-loops compared to suppressive R-loops of these regions including 5’ end, body, and 3’ end in those cell lines. For example, in Hela-S3 cell line, there are 8,746 5’ end proliferative R-loops, 4,933 body proliferative R-loops, and 6,304 3’end proliferative R-loops, but the number for 5’end suppressive R-loops, body suppressive R-loop, 3’end suppressive R-loops is only 2,880, 3,862, 4,575 respectively (Supplementary Table S1). This may indicate aberrant accumulation of R-loops is the main cause of cancer development and progression. To note, the degree of aberrant R-loop varies from cell line to cell line as well. For example, there exists a more overwhelming degree of aberrant R-loops in Hela than U87, U2OS, and K562, suggesting aberrant R-loops may play a cell line-specific role in biological processes.

### The architecture of Deep R-looper Discriminant

Here, we proposed a deep neural network-based framework named Deep R-looper Discriminant, for extracting features automatically from epigenetic marks in genome bins around TSS and TTS and identifying aberrant R-loops against normal R-loops. Figure 2A illustrates the architecture of the Deep R-looper Discriminant. Basically speaking, the epigenetic marks (DNase, RNAPII, H3K4me3, H3K27ac, H3K9ac, H3K27me3, H3K9me3, H3K4me1, ATAC) of 80 bins around TSS and TTS altogether are taken as an input, the input is input as 9*160 matrix. The first layer is a linear layer that flatten the initial input matrix. After that, the reconstructed input is fed into one continuous bidirectional long short-term memory layer (BLSTM) and the layer integrates information from forward and backward layers and captures intrinsic features in a supervised manner, hence improving the final predictive performance by keeping location information[44]. Afterward, the concatenated information from the BLSTM layer propagates to a linear layer, and is converted to 1D vector, and then is fed into three consecutive fully-connected (FC) layers where batch normalization is applied to extract information further and prepare it for the fourth fully-connected layer consisting of one unit to make a final prediction, where the sigmoid activation function is applied to achieve the classification task appropriately. Batch normalization serves to standardize the mean and variance of each units to stabilize learning, and is proved to facilitate and coordinate the update of multiple layers successfully[45].

**Figure 2.**
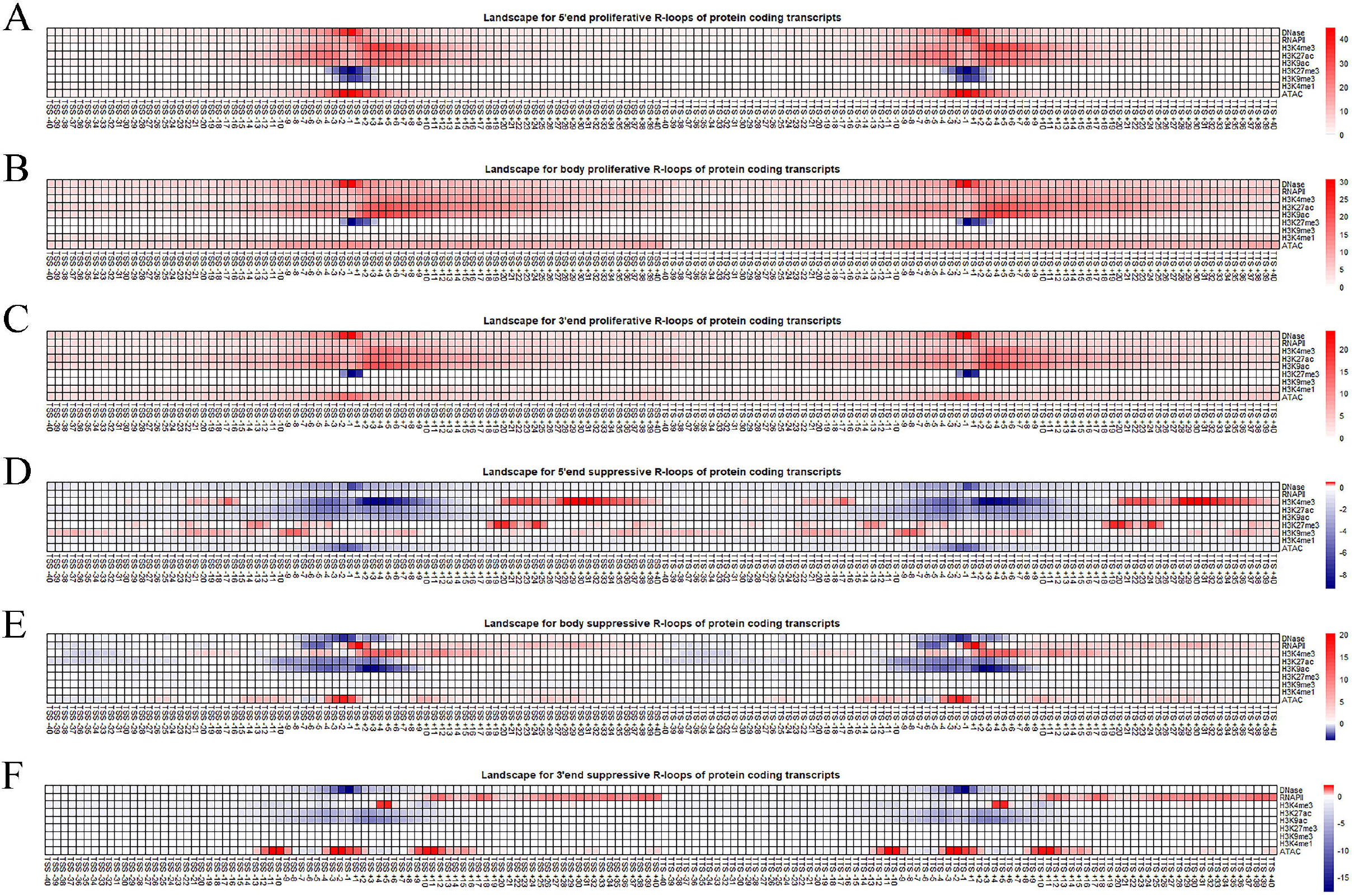
The framework of Deep R-looper Discriminant and its application in identifying aberrant R-loops. (A) The architecture of Deep R-looper Discriminant. (B) Predictive performance evaluation for Deep R-looper Discriminant and baselines when identifying aberrant R-loops against normal R-loops in protein-coding transcripts. (C) Predictive performance evaluation for Deep R-looper Discriminant and baselines when identifying aberrant R-loops against normal R-loops in non-coding transcripts.

Here, we adopted multiple learning techniques to facilitate model training and make the framework more accurate and robust. Dropout is applied to the intermediate between FC layers to overcome overfitting. Dropout is thought as the most common regularization method and a key component to training complicated deep neural network models, and it can set some neurons in the network to “dropped out” state during training in each iteration, which can average multiple networks during training and make the final trained model more generalized in essence[46]. Besides, as an effective regularization method, early stopping is also adopted to stop training with patience once the model performance was no longer improved on the validation dataset to the aim of avoiding overfitting. L1 norm and L2 norm are also combined for regularizing the parameter values in the FC layers. To find out the best parameter combination, the initialized value of each parameter is sampled from a uniform or normal distribution randomly in the BLSTM layer, and the FC layers. Lastly, for obtaining better local minima of parameters, adadelta learning rate method is adopted for gradient descent, which was proved to be robust to noisy gradient information, different model architecture, and diverse data modalities[47].

In the Deep R-looper Discriminant framework, several hyperparameters needed to be searched for the best fit during the training process, including dropout rate, batch size, units in the FC layers, etc. The predictive performance of hyperparameter combinations was evaluated based on 5-fold cross-validations. To get the best combination of hyperparameters in the high-dimensional hyperparameter space effectively, Bayesian optimization was adopted in the training process. During the training process, as iteration grew, and the posterior distribution of the model’s cost function improved, the Bayesian optimization algorithm could further explore hyperparameter space which was worth exploring automatically, and sought the best hyperparameter combination as soon as possible.

### The evaluation of the contributions of spatial multi-omics to aberrant R-loops

To identify those key spatial epigenetic marks that may contribute to aberrant R-loops, including proliferative R-loops and suppressive R-loops of 5’ end, body and 3’ end of transcripts, we adopted the permutation test method based on ELI5 to compute feature importance derived from the finalized Deep R-looper Discriminant models trained on specific datasets respectively by measuring how importance score decreases when each spatial epigenetic mark is not available. This method randomly shuffles each input feature and computes the change in the model’s predictive performance. Therefore, we finally could get key spatial epigenetic marks that are tightly related to aberrant R-loops.

### Modelling the relationship between epigenetic marks and aberrant R-loops

We also constructed multiple machine learning models to serve as baselines and illuminate the relationship between epigenetic marks and aberrant R-loops. The models include logistic regression (LR), k-nearest neighbors (KNN), naive bayes (NB), random forests (RF), and linear discriminant analysis (LDA). All these models are applied to the below two tasks: identifying aberrant R-loops against normal R-loops; classifying the class of aberrant R-loops.

For the task to identify aberrant R-loops against normal R-loops and the task to classify the class of aberrant R-loops of the same region, we evaluated the predictive performance of the models based on the F1-score considering there are unbalanced datasets.

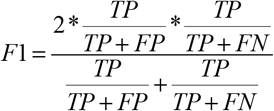

To evaluate the predictive performance of the Deep R-looper Discriminant comprehensively, we compared the model to the abovementioned baselines, including LR, LDA, KNN, NB and RF after grid search optimization in the aspect of F1 score based on 5-fold cross-validation. The detailed hyperparameter space of baselines modelling methods and Deep R-looper Discriminant are shown in Supplementary Table S2.

### Deep R-looper Discriminant achieves robust performance for identifying aberrant R-loops against normal R-loops using epigenetic marks

The Deep R-looper Discriminant is a competent framework to identify aberrant R-loops against housekeeping R-loops based on epigenetic marks across cell lines. After tuning the hyperparameter via Bayesian optimization and got the optimized configurations of the Deep R-looper Discriminant, we also adopted GridSearch CV to automate the tuning of hyperparameters for these baseline models and finally got optimized LR, KNN, LDA, RF, and NB. To evaluate Deep R-looper Discriminant comprehensively, we compared its performance against baselines in the aspect of F1 score.

Obviously, Deep R-looper Discriminant achieved the best performance regarding F1-score among these prediction models across cell lines in protein coding transcripts. For example, in K562 cell line, Deep R-looper Discriminant achieved F1 at 0.93±0.078 (standard deviation, SD); however, LR got 0.24±0.046 (SD), LDA got 0.28±0.025 (SD), KNN got 0.40±0.034 (SD), NB got 0.06±0.007 (SD), and RF got 0.36±0.033 (SD). The superior predictive performance of Deep R-looper Discriminant is also observed in Hela, U2OS, and U87 cell lines (Figure 2B).

When it comes to identifying aberrant R-loops against normal R-loops across cell lines in non coding transcripts, Deep R-looper Discriminant also showed more promising predictive performance than baselines. For example, in U2OS cell line, Deep R-looper Discriminant achieved F1 at 0.91±0.041 (SD); however, LR got 0.36±0.038 (SD), LDA got 0.46±0.017 (SD), KNN got 0.43±0.028 (SD), NB got 0.21±0.011 (SD), and RF got 0.40±0.041 (SD). The similar trend is also observed in Hela, K562, and U87 cell lines (Figure 2C). The detailed predictive performance in this part is shown in Supplemental Table 3.

Taken all altogether, these results suggest although the predictive performance of Deep R-looper Discriminant varies when identifying aberrant R-loops against normal R-loops across cell lines and the types of transcripts, the framework is a great fit for identifying aberrant R-loops in protein coding transcripts and even non coding transcripts and is able to capture conserved patterns distinguishing aberrant R-loops from normal R-loops and could be generalized across cell lines.

### Deep R-looper Discriminant achieves robust performance for classifying aberrant R-loops using epigenetic marks

Furthermore, we explored whether Deep R-looper Discriminant was able to distinguish between proliferative and suppressive R-loops at the same regions, including 5’ end, body, 3’ end. Similarly, after tuning the hyperparameters, we got multiple optimized prediction models to distinguish between proliferative and suppressive R-loops at 5’ end, body, 3’ end across cell lines and the types of transcripts respectively based on F1 score.

Again, the optimized Deep R-looper Discriminant achieved the best performance regarding F1-score among these prediction models across cell lines in protein coding transcripts. For example, for protein coding transcripts, Deep R-looper Discriminant achieved F1 at 0.98±0.005 (SD) for aberrant 5’ end R-loops of U87, 1.00±0.002 (SD) for aberrant body R-loops of U87 and 1.00±0.003 (SD) for aberrant 3’ end R-loops of U87; however, LR got 0.88±0.003 (SD), 0.91±0.006 (SD), 0.91±0.005 (SD), LDA got 0.89±0.008 (SD), 0.77±0.041 (SD), 0.75±0.019 (SD), KNN got 0.81±0.005 (SD), 0.87±0.007 (SD), 0.86±0.003 (SD), NB got 0.80±0.012 (SD), 0.91±0.008 (SD), 0.90±0.009 (SD), and RF got 0.88±0.005 (SD), 0.90±0.005 (SD), 0.90±0.001 (SD) respectively. The superior predictive performance of Deep R-looper Discriminant is also observed in Hela, U2OS, and U87 cell lines (Figure 3A, 3C and 3E).

**Figure 3.**
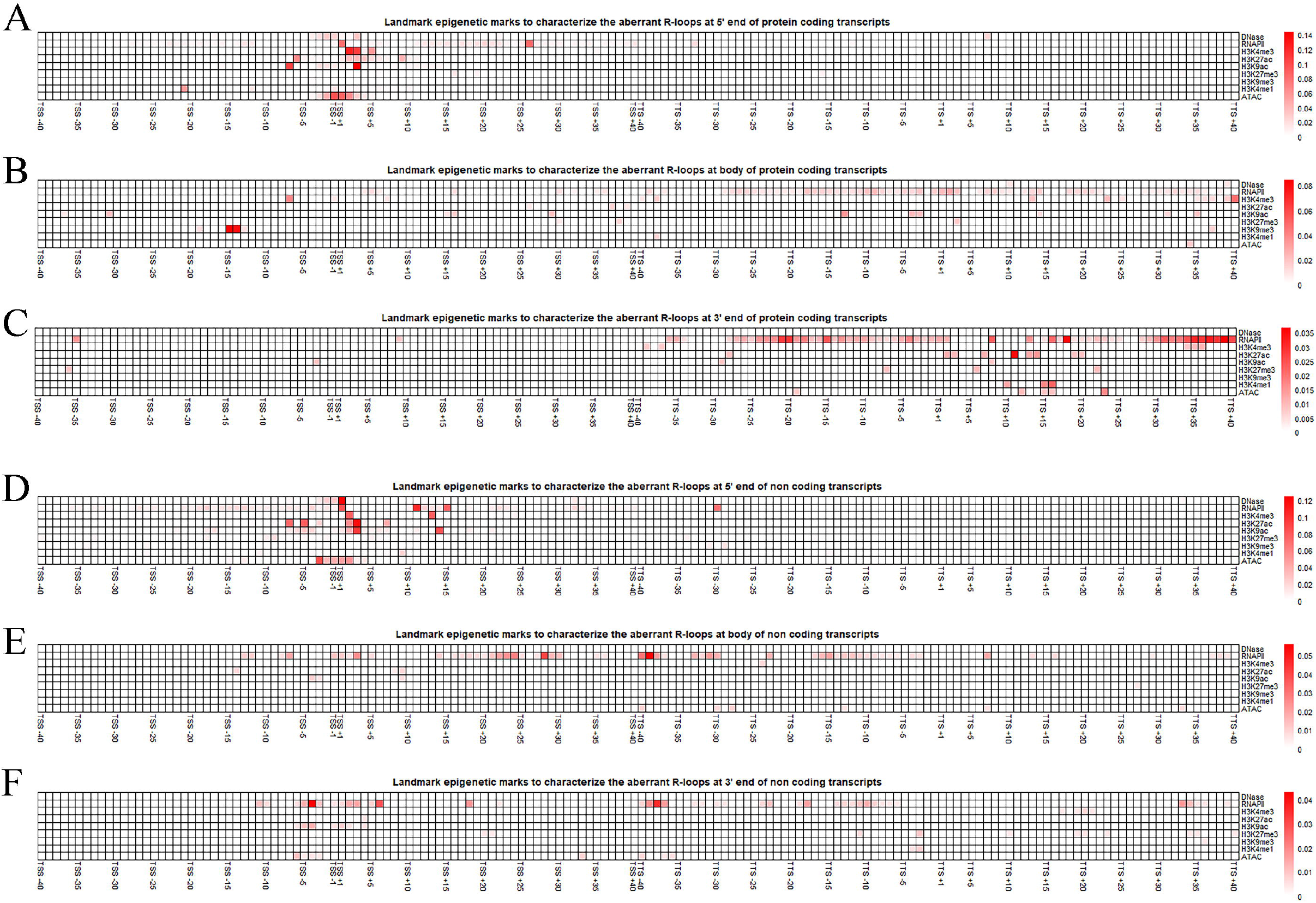
Deep R-looper Discriminant’s application in identifying aberrant 5’ end, body, 3’ end R-loops. (A) Predictive performance evaluation for Deep R-looper Discriminant and baselines when classifying aberrant 5’ end R-loops in protein-coding transcripts. (B) Predictive performance evaluation for Deep R-looper Discriminant and baselines when classifying aberrant 5’ end R-loops in non-coding transcripts. (C) Predictive performance evaluation for Deep R-looper Discriminant and baselines when classifying aberrant body R-loops in protein-coding transcripts. (D) Predictive performance evaluation for Deep R-looper Discriminant and baselines when classifying aberrant body R-loops in non-coding transcripts. (E) Predictive performance evaluation for Deep R-looper Discriminant and baselines when classifying aberrant 3’ end R-loops in protein-coding transcripts. (F) Predictive performance evaluation for Deep R-looper Discriminant and baselines when classifying aberrant 3’ end R-loops in non-coding transcripts.

When it comes to non coding transcripts, Deep R-looper Discriminant also showed more promising predictive performance than baselines. Deep R-looper Discriminant achieved F1 at 0.99±0.009 (SD) for aberrant 5’ end R-loops of U2OS, 0.97±0.022 (SD) for aberrant body R-loops of U2OS and 0.95±0.030 (SD) for aberrant 3’ end R-loops of U2OS; however, LR got 0.86±0.019 (SD), 0.80±0.043 (SD), 0.77±0.050 (SD), LDA got 0.64±0.016 (SD), 0.54±0.076 (SD), 0.60±0.023 (SD), KNN got 0.78±0.010 (SD), 0.77±0.047 (SD), 0.68±0.055 (SD), NB got 0.88±0.012 (SD), 0.75±0.053 (SD), 0.72±0.040 (SD), and RF got 0.87±0.017 (SD), 0.83±0.029 (SD), 0.76±0.032 (SD) respectively. The superior predictive performance of Deep R-looper Discriminant is also observed in Hela, U2OS, and U87 cell lines (Figure 3B, 3D and 3F). The detailed predictive performance in this part is shown in Supplemental Table 3.

The robust predictive performance of Deep R-looper Discriminant is proved to be successful in these scenarios, suggesting Deep R-looper Discriminant is able to capture and learn the complex and intrinsic relationship between proliferative R-loop and suppressive R-loop effectively.

### The landscape of Epigenetic marks in aberrant R-loops

It has been reported that R-loop represents a kind of epigenetic information and is accompanied by a characteristic set of histone modifications[3, 5, 48-50]. For instance, promoter R-loop associates with transcription, so active transcription marks such as H3K4me1 usually occur at promoter regions with R-loop[3, 5]. R-loop also serves in chromatin decondensation and opens the chromatin structure, therefore it positively associates with DNase I hyper accessibility as well[3, 51-54]. Besides, there is a strong correlation between RNAP II stalling and R-loop, considering either of them may be promoted by the other and both participate in the regulation of transcription and expression[3, 5, 48-50].

To characterize the epigenetic marks of those aberrant R-loops comprehensively, we depicted the landscape of DNase, RNAPII, H3K4me3, H3K27ac, H3K9ac, H3K27me3, H3K9me3, H3K4me1, ATAC) of 80 bins around TSS and TTS by comparing aberrant R-loops to normal R-loops.

For proliferative R-loops of 5’ end, body, 3’ end of protein coding transcripts (Figure 4A, 4B, and 4C), although they have similar epigenetic features in space where DNase, RNAPII, H3K4me3, H3K27ac, H3K9ac, H3K4me1, and ATAC show a widespread increased trend, they show subtle difference where there is the peakiest distribution for 5’ end proliferative R-loops but the distribution is more dispersive for 3’ end and body proliferative R-loops (Figure 4A, 4B, and 4C). Besides, 5’ end proliferative R-loops usually a decreased H3K27me3 and H3K9me3 in bins adjacent to TSS and TTS, but body proliferative R-loops and 3’ end proliferative R-loops only have a decreased H3K27me3 in bins adjacent to TSS and TTS.

**Figure 4.**
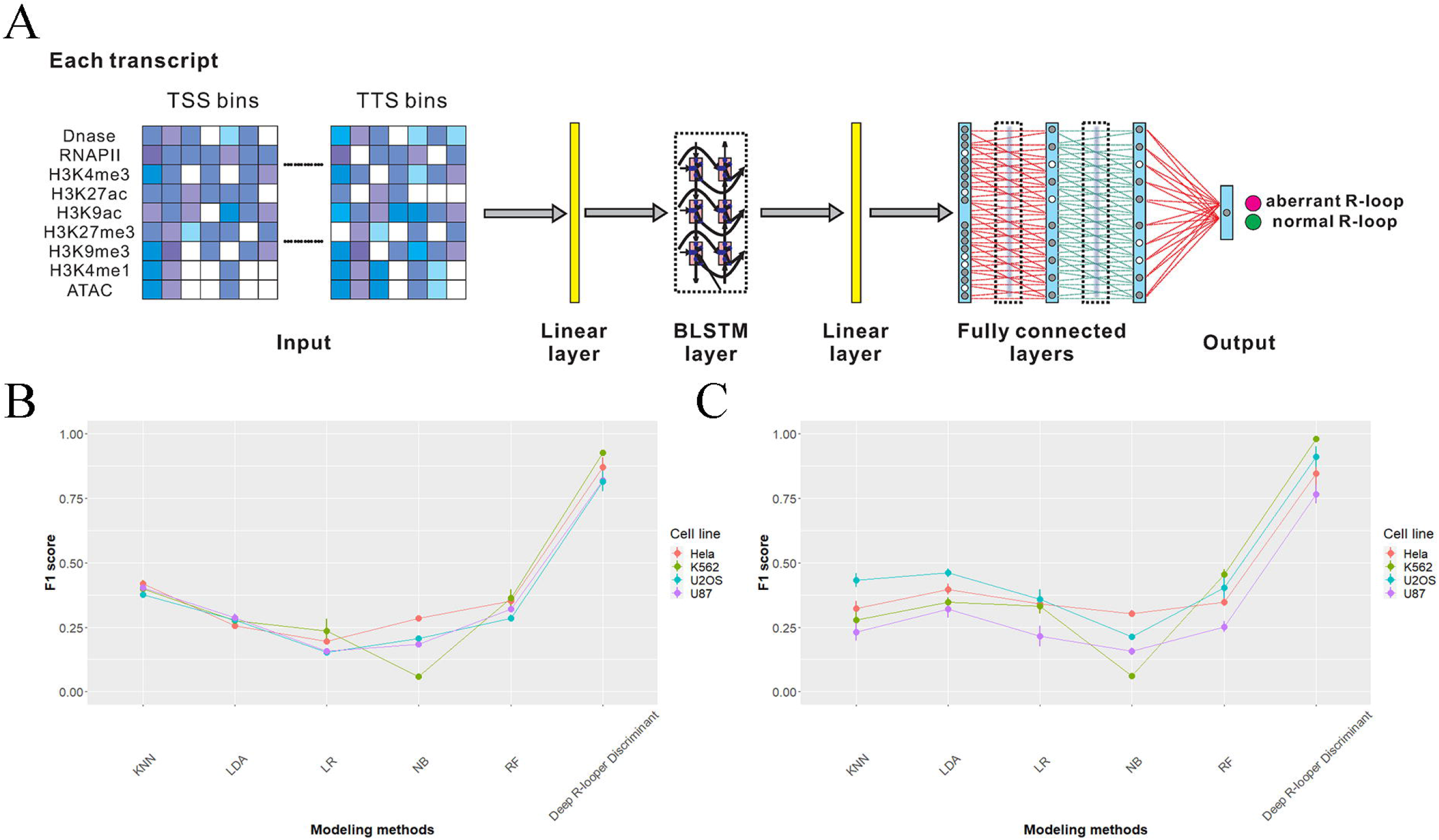
Aberrant R-loops landscape. (A) Landscape for 5’ end proliferative R-loops of protein coding transcripts. (B) Landscape for body proliferative R-loops of protein coding transcripts. (C) Landscape for 3’ end proliferative R-loops of protein coding transcripts. (D) Landscape for 5’ end proliferative R-loops of non coding transcripts. (E) Landscape for body proliferative R-loops of non coding transcripts. (F) Landscape for 3’ end proliferative R-loops of non coding transcripts.

In contrast, suppressive R-loops of 5’ end, body, 3’ end of protein coding transcripts show complicated epigenetic features in genome bins. Generally speaking, although they all have two discrete regions adjacent to TSS and TTS where DNase, H3K27ac, and H3K9ac show a decreased trend, 5’ end suppressive R-loops also show the decrease of H3K4me3, ATAC in these two regions. To note, 5’ end suppressive R-loops show a strong decrease of ATAC in regions adjacent to TSS and TTS. However, body suppressive R-loops and 3’ end suppressive R-loops show a strong increase of ATAC in the regions (Figure 4D, 4E, and 4F).

When it comes to proliferative R-loops in non coding transcripts, they stand in sharp contrast to those in protein coding transcripts (Supplementary Figure S1). For example, proliferative R-loops of 5’ end, body, 3’ end in non coding transcripts show a widespread decrease in H3K27me3, H3K9me3, and H3K4me1, and meanwhile, these R-loops possess multiple discrete decrease regions of DNase, and H3K4me3. To note, suppressive R-loops in non coding transcripts stand in sharp contrast to those in protein coding transcripts as well (Supplementary Figure S1). All these differences suggest aberrant R-loops may be associated with the transcription process.

### Landmark epigenetic modifications contribute to differentiation between proliferative and suppressive R-loops in a location-specific manner

Although epigenetic marks are associated with aberrant R-loop cooperatively, their contributions may be not identical in genome backgrounds. To further investigate which epigenetic modification contributes to the differentiation between proliferative and suppressive R-loops, we, therefore, computed and evaluated the relative contribution of spatial epigenetic marks to aberrant R-loops across cell lines and the types of transcripts. More specifically, after optimizing the configurations of Deep R-looper Discriminant, we applied the optimized models to corresponding datasets and got importance score of spatial multi-omics features based on permutation tests. We finally evaluated all epigenetic marks across 80 bins around TSS and TTS based on their contribution score to the formation of 5’ end aberrant R-loops, body aberrant R-loops, 3’ end aberrant R-loops respectively.

For aberrant R-loops at the 5’ end of protein coding transcripts, it shows that H3K4me3, ATAC, H3K9ac, H3K27ac, and RNAP II in TSS proximal regions are landmark modifications to the differentiation between 5’ end proliferative and suppressive R-loops. Besides, RNAPII in TSS downstream distal region (TSS +26) and H3K4me1 in TSS upstream distal region (TSS -21) exert influence as well (Figure 5A). However, the epigenetic marks residing in TTS didn’t seem to exert an influence on these two kinds of R-loops. As for aberrant R-loops at the body of protein coding transcripts, RNAPII, H3K4me3, H3K9ac, and H3K9me3 in both TSS and TTS regions contribute more to the differentiation of aberrant body R-loops than DNase, H3K27ac, H3K27me3, H3K4me1, and ATAC. Interestingly, the contribution of DNase, H3K4me1, and ATAC is only confined in TTS regions and that of H3K27ac is only confined in TSS regions (Figure 5B). In contrast to R-loops at 5’ end of protein coding transcripts, the contribution of epigenetic marks to the 3’ end aberrant R-loops is mainly from TTS regions, including RNAPII, H3K4me3, H3K27ac, H3K9ac, H3K27me3, H3K4me1, and ATAC (Figure 5C).

**Figure 5.**
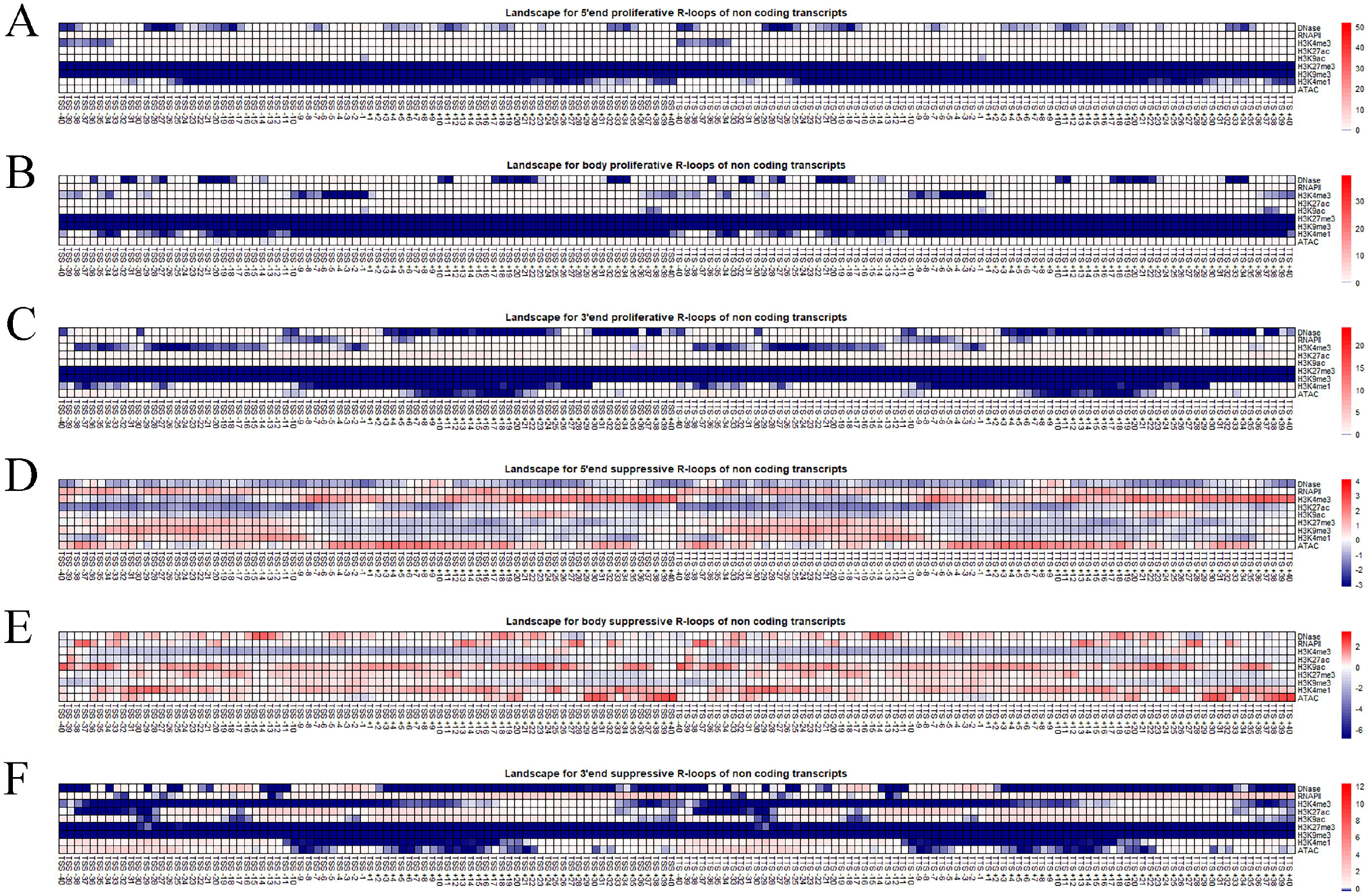
Landmark epigenetic marks to characterize the aberrant R-loops of protein coding transcripts. (A) Landmark epigenetic marks to characterize the proliferative R-loops at 5’ end of protein coding transcripts. (B) Landmark epigenetic marks to characterize the proliferative R-loops at body of protein coding transcripts. (C) Landmark epigenetic marks to characterize the proliferative R-loops at 3’ end of protein coding transcripts. (D) Landmark epigenetic marks to characterize the suppressive R-loops at 5’ end of protein coding transcripts. (E) Landmark epigenetic marks to characterize the suppressive R-loops at body of protein coding transcripts. (F) Landmark epigenetic marks to characterize the suppressive R-loops at 3’ end of protein coding transcripts.

Interestingly, the contribution pattern of epigenetic marks to aberrant R-loops in non coding transcripts is kind of similar to that in protein-coding transcripts, suggesting the differentiation may be transcription-independent (Figure 5D, E, F).

Taken all altogether, these results suggest the epigenetic marks may play a cis role, and local epigenetic features adjacent to TSS and TTS regions may be the key to the formation of proximal aberrant R-loops.

### Target genes regulated by R-loop paly a cell line-specific role

Integrated with publicly available RNA expression datasets and R-loops profiling datasets across cell lines, we identified target genes regulated directly by R-loops using BETA[55]. Although there are 983, 217, 118, and 160 up-regulated target genes by R-loops, none of them is in common (Figure 6A). Similarly, there are 1,164, 160, 103, and 250 down-regulated target genes regulated by R-loops, and none of them is in common as well (Figure 6D). These finding suggest that these target genes regulated by R-loops may influence in a context specific way. The detailed target genes regulated by R-loops are shown in Supplemental Table 4.

**Figure 6.**
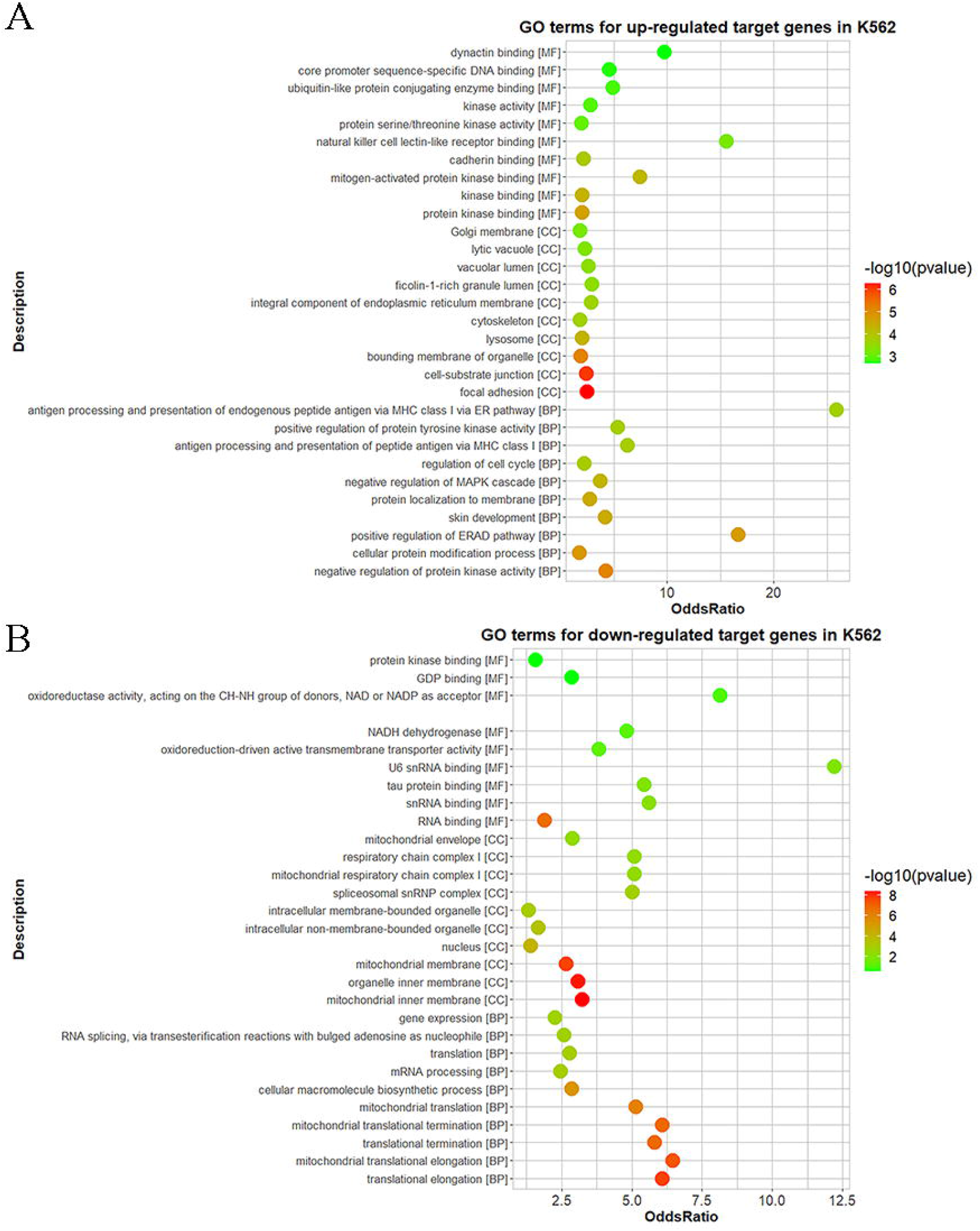
Target genes that tare regulated by R-loops across cell lines. (A) Up-regulated target genes regulated by R-loops across cell lines. (B) Pathway analysis for target genes regulated by R-loops in K562. (C) Pathway analysis for target genes regulated by R-loops in U2OS. (D) down-regulated target genes regulated by R-loops across cell lines. (E) Pathway analysis for target genes regulated by R-loops in Hela. (F) Pathway analysis for target genes regulated by R-loops in U87.

Furthermore, we carried out pathway analysis for those target genes regulated by R-loops. For example, in K562, for up-regulated target genes, they are involved in regulation of actin cytoskeleton, focal adhesion; for down-regulated target genes, they involve in DNA replication, Kit receptor signalling pathway and so on (Figure 6B). Similarly, quite a few context-specific pathways are enriched by these target genes regulated R-loops in U2OS, Hela and U87 cell lines, suggesting R-loop participate in cell line specific biological pathways by regulating target genes (Figure 6D, 6E and 6F). Further GO enrichment analysis also support this cell line specific regulation by R-loops across cell lines (Supplementary Figure S2, S3, S4, and S5).

Besides, after motif analysis for target regions by R-loops, a cell line specific transcription factor (TF) landscape regulated by R-loops are observed (Supplementary Figure S6 and S7). There are several shared up-regulated TFs by R-loops across cell lines, including CUX1, YY1, POU3F1, POU3F2, POU3F3, FOXO3 and so on. As for common down-regulated TFs by R-loops across cell lines, they include some YY1, FOXO3, POU3F2, POU3F3 and so on. To note, YY1, POU3F2/3 and FOXO3 are found to be common TFs that can be up-regulated and down-regulated by R-loops across cell lines, suggesting these TFs are key factors that participate in R-loop regulations. The detailed TFs regulated by R-loops are shown in Supplemental Table 5.

## DISCUSSION

Abnormal R-loops have been found to the cause of multiple human diseases, including cancers and neurological disorders. However, there is a lacking in defining what is “aberrant” R-loops. Here, we are first to collect loads of publicly available R-loop profiling data and define aberrant R-loops at the whole genome level systematically, and we further subclass those aberrant R-loops into six classes, including 5’ end proliferative R-loop, body proliferative R-loop, 3’ end proliferative R-loop, 5’ end suppressive R-loop, body suppressive R-loop, and 3’ end suppressive R-loop. It shows aberrant R-loops are prevalent across cell lines and also the degree of aberrant R-loops varies from cell line to cell line, indicating aberrant R-loops may play a cell line-specific role.

Considering aberrant R-loops only account for a minority of transcripts at the whole genome level, how to identify aberrant R-loops against loads of normal R-loops is more and more necessary. Here we proposed an original deep neural network-based framework named Deep R-looper Discriminant for the task. By comparing with other benchmark models, the framework achieved a robust predictive performance in identifying aberrant R-loops against other normal R-loops. Furthermore, considering there are diverse aberrant R-loops and different subclasses of aberrant R-loops may show distinctive characteristics, therefore we next adjusted the framework to make it available to subclassify the aberrant R-loops into the six abovementioned subclasses. Again, our framework achieves a robust predictive performance in subclassifying the aberrant R-loops. Based on the same architecture, the models are capable of achieving predictive performance in identifying aberrant R-loops against normal R-loops and subclassifying aberrant R-loops into subclasses across cell lines, indicating our framework is a robust framework to be applied to unknown cell line and also there is cell line specificity of aberrant R-loops in organism.

By comparing normal R-loops with the six subclasses of aberrant R-loops, we found the distinctive spatial histone landscape for these aberrant R-loops. For example, the characteristic of 5’ end proliferative R-loops is the prevalent decrease of H3K27me3 in bins around TSSs and a minor decrease of H3K9me3 around bins close to TSSs. Besides, we evaluated the histone marks in each bin according to their contribution score to the formation of the aberrant R-loops and then determined the landmark histone marks for the formation of aberrant R-loops.

By integrating available omics data across cell lines, we identified regulatory target genes by R-loop across cell lines, including up-regulated targets and down-regulated targets. By comparing up-regulated genes and down-regulated genes across cell lines, we found most of them are exclusive in context specific way, therefore we infer that R-loops exert a cell line specific biological pathways and functions. Furthermore, we identified key TFs that are regulated in R-loops across cell lines, some of them are conserved such as POU3F2/3 and FOXO3.

---

**SFigure 1.**
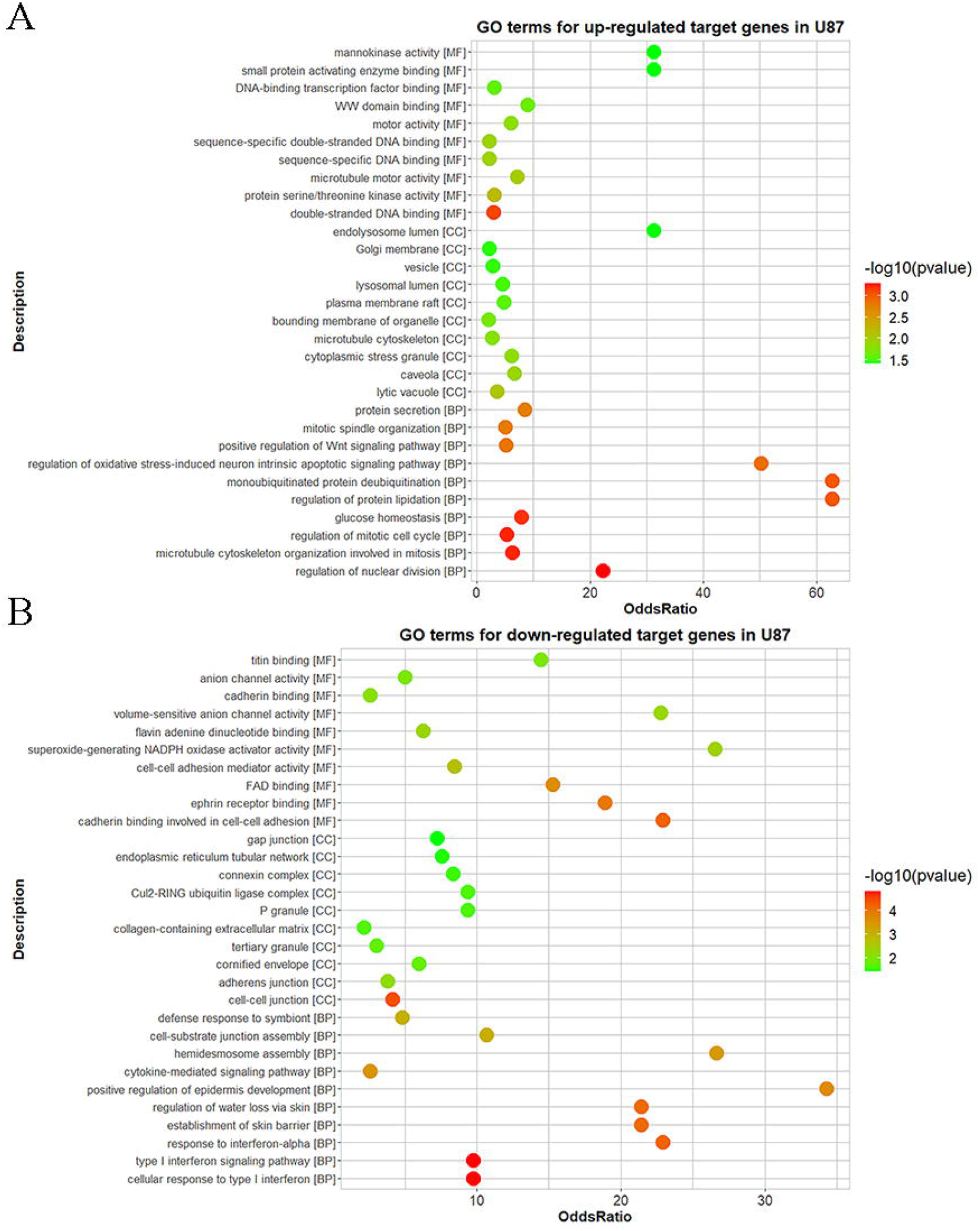
Landmark epigenetic marks to characterize the aberrant R-loops of non coding transcripts. (A) Landmark epigenetic marks to characterize the proliferative R-loops at 5’ end of non coding transcripts. (B) Landmark epigenetic marks to characterize the proliferative R-loops at body of non coding transcripts. (C) Landmark epigenetic marks to characterize the proliferative R-loops at 3’ end of non coding transcripts. (D) Landmark epigenetic marks to characterize the suppressive R-loops at 5’ end of non coding transcripts. (E) Landmark epigenetic marks to characterize the suppressive R-loops at body of non coding transcripts. (F) Landmark epigenetic marks to characterize the suppressive R-loops at 3’ end of non coding transcripts.

**SFigure 2.**
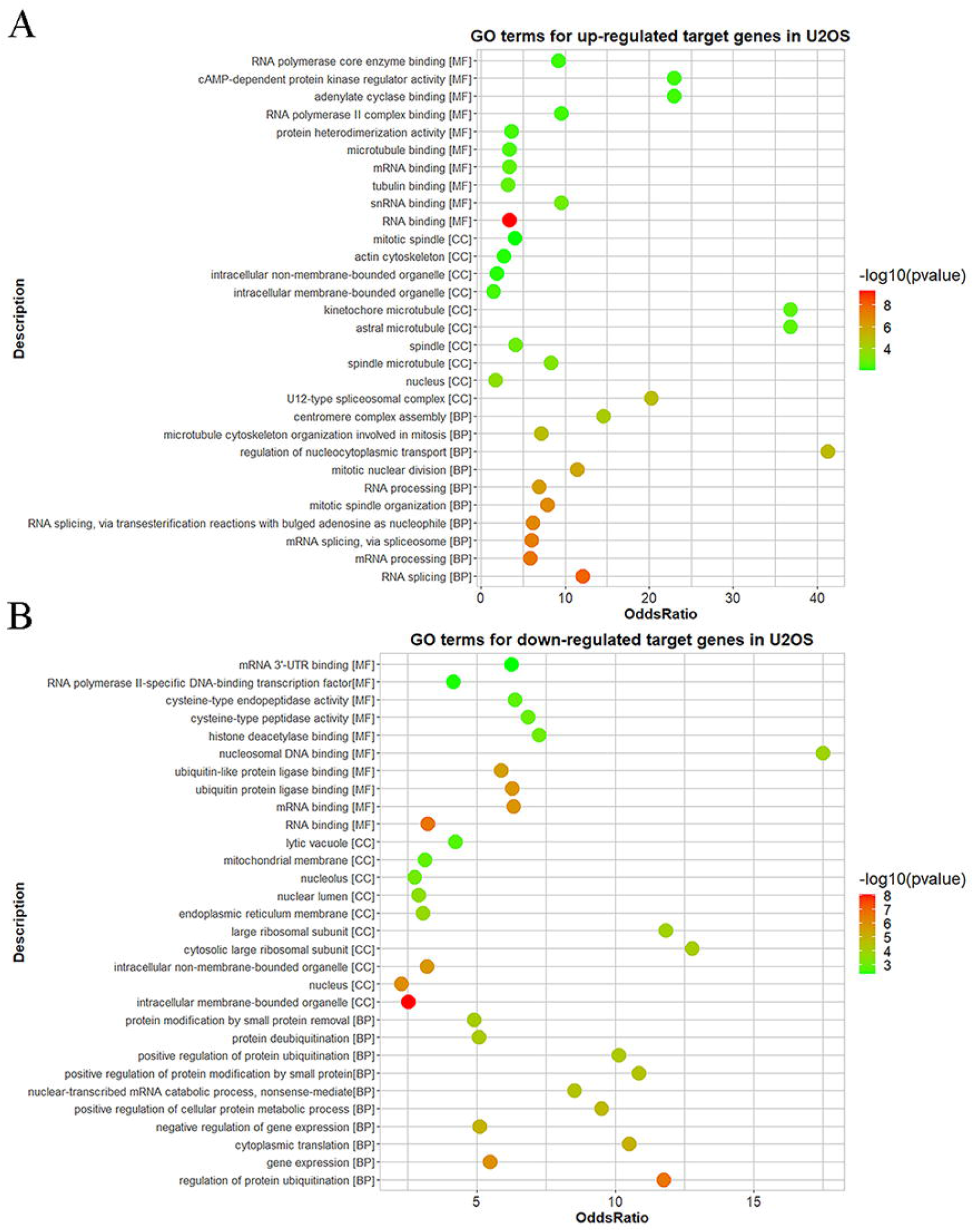
GO terms for regulated target genes by R-loop in K562. (A) GO terms for up-regulated target genes by R-loops in K562 (B) GO terms for down-regulated target genes by R-loops in K562

**SFigure 3.**
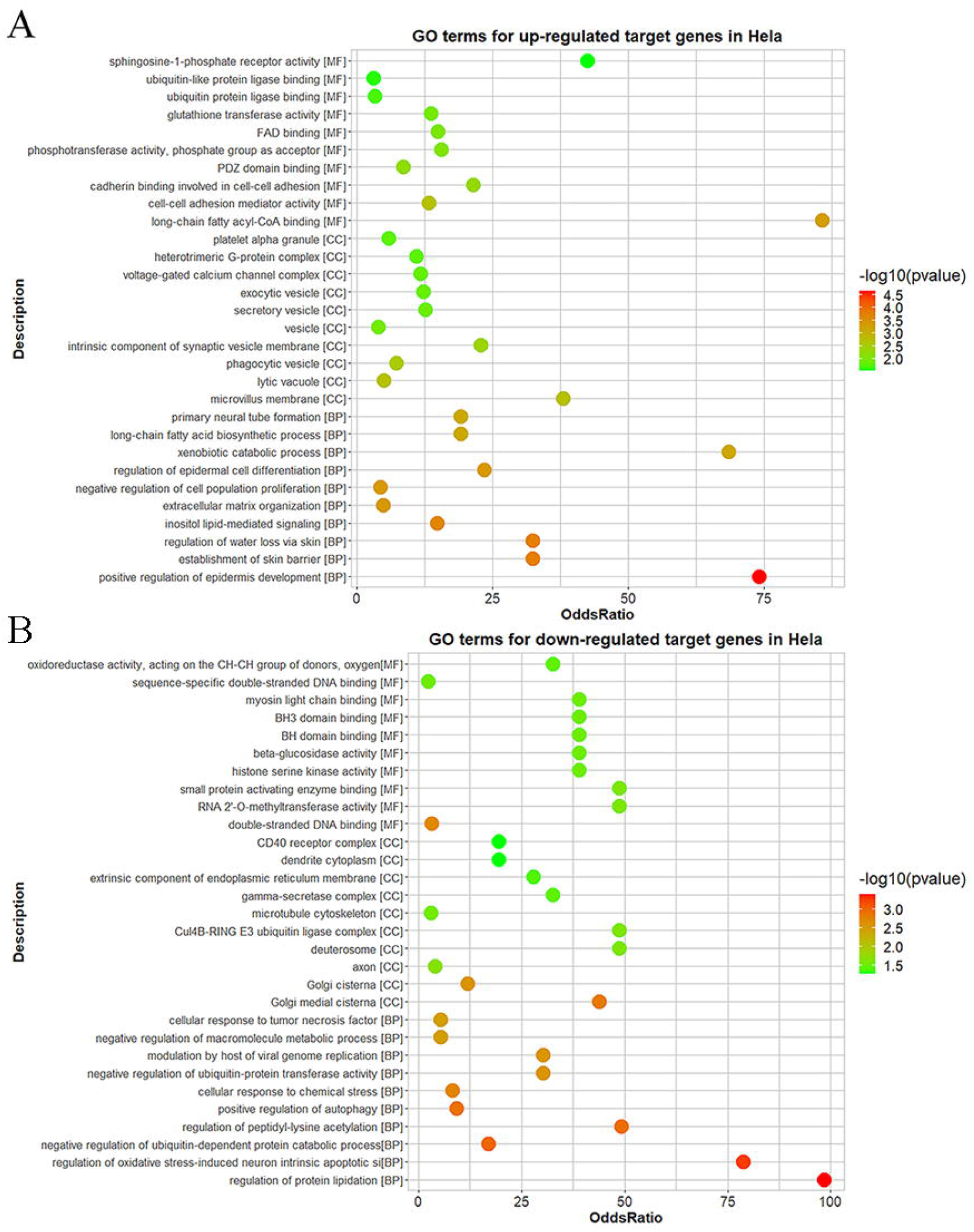
GO terms for regulated target genes by R-loop in U87. (A) GO terms for up-regulated target genes by R-loops in U87 (B) GO terms for down-regulated target genes by R-loops in U87

**SFigure 4.**
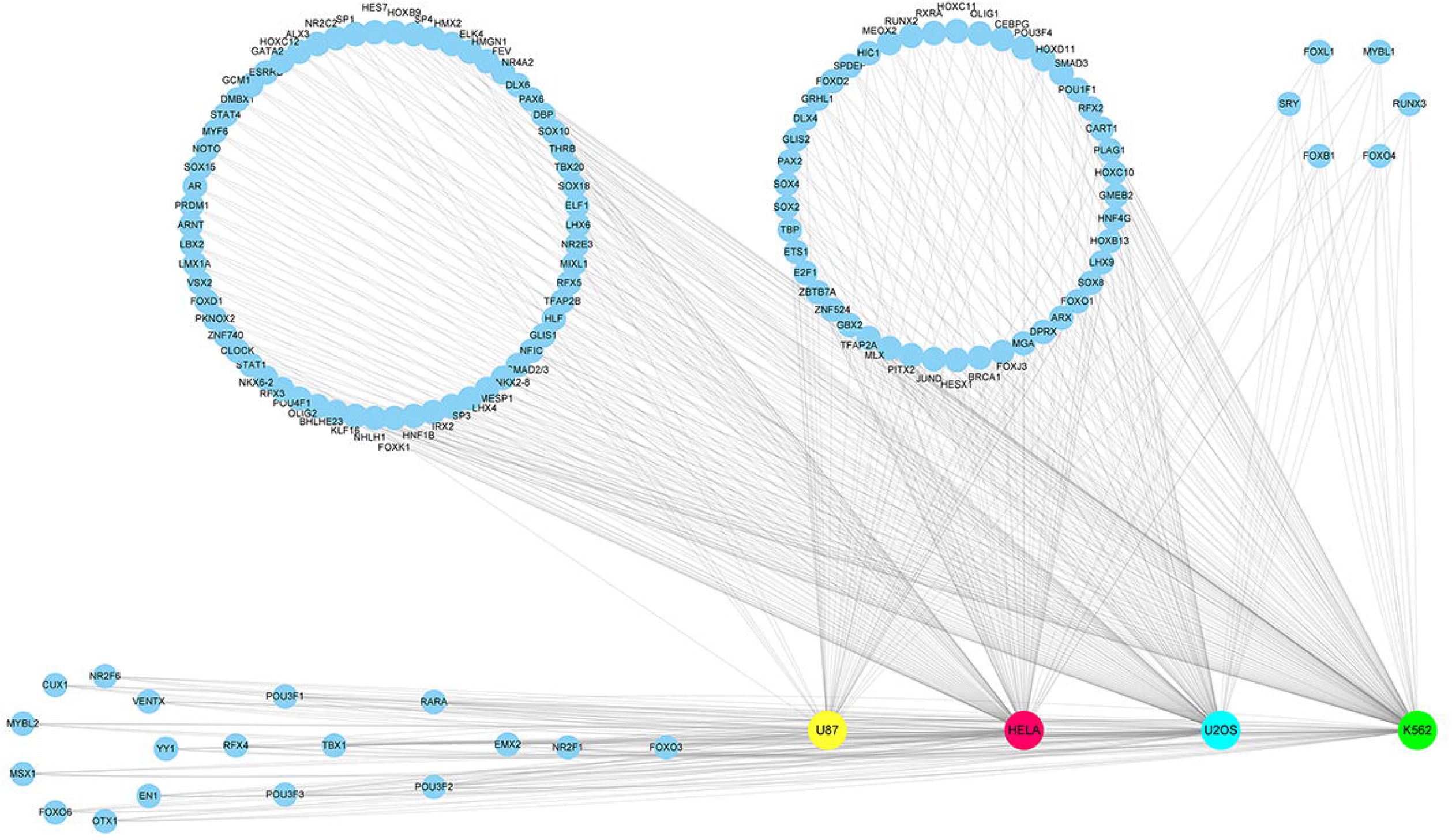
GO terms for regulated target genes by R-loop in U2OS. (A) GO terms for up-regulated target genes by R-loops in U2OS (B) GO terms for down-regulated target genes by R-loops in U2OS

**SFigure 5.**
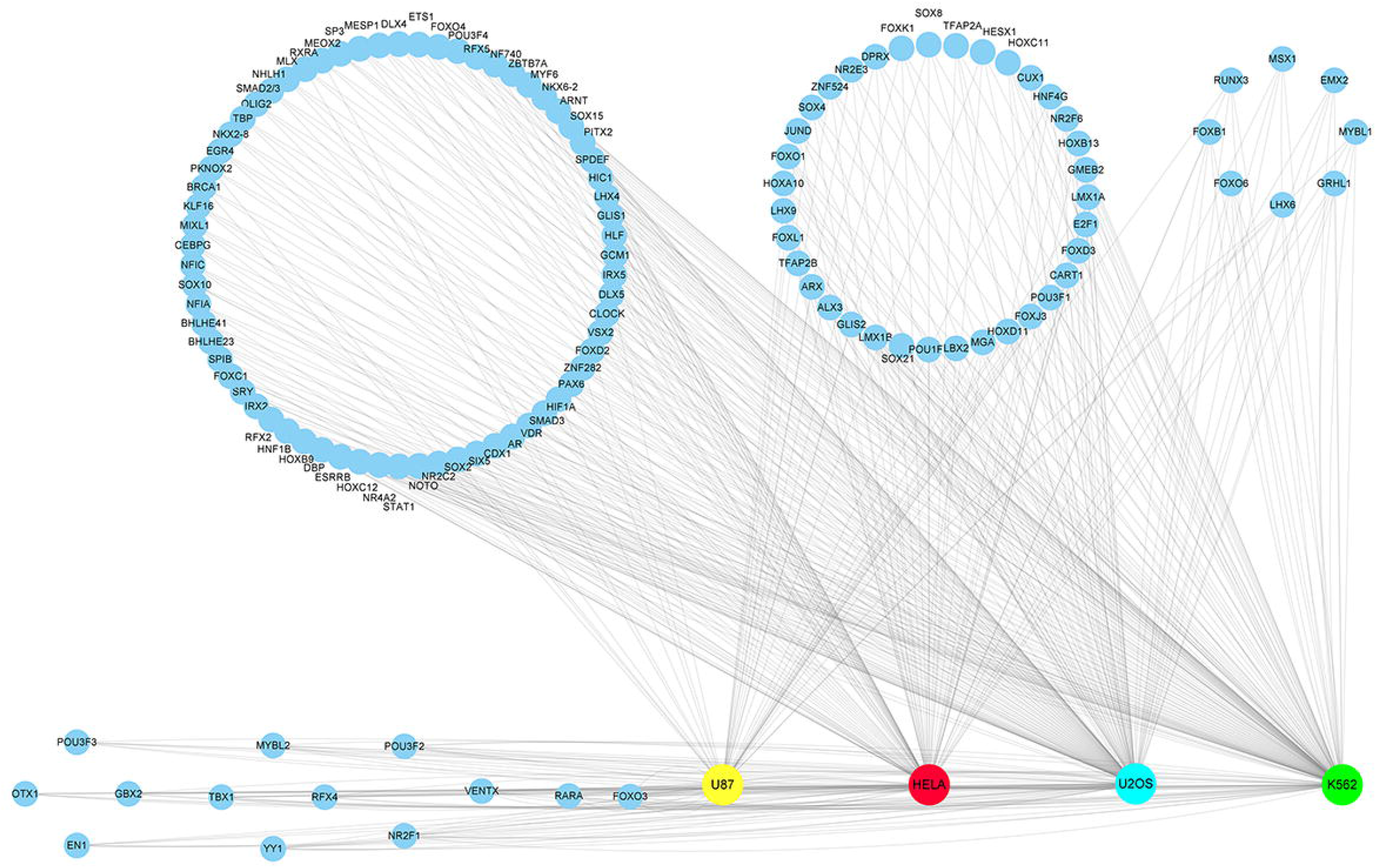
GO terms for regulated target genes by R-loop in Hela. (A) GO terms for up-regulated target genes by R-loops in Hela (B) GO terms for down-regulated target genes by R-loops in Hela

**SFigure 6.**
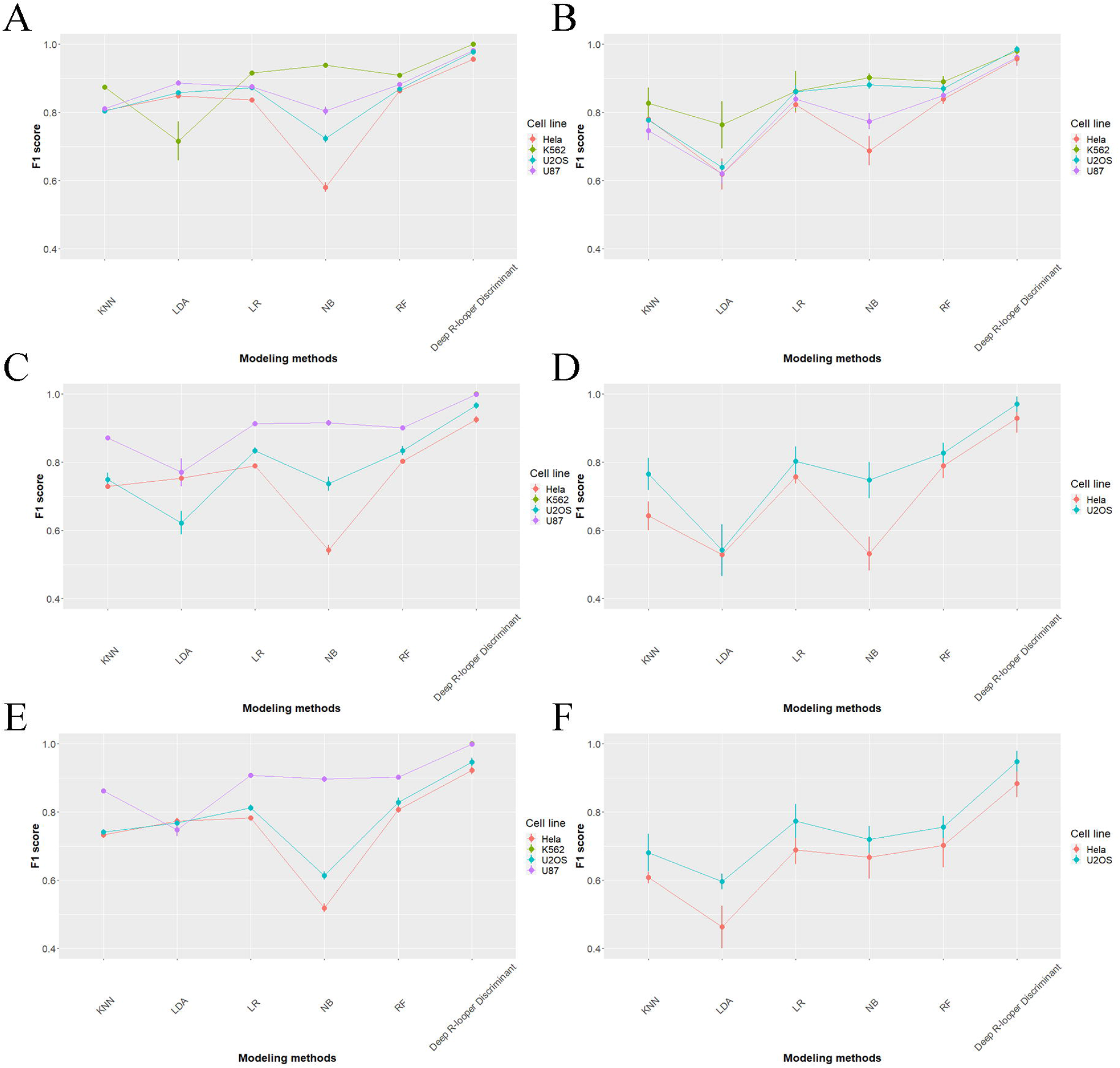
Transcription factors that are up-regulated by R-loops across cell lines.

**SFigure 7.**
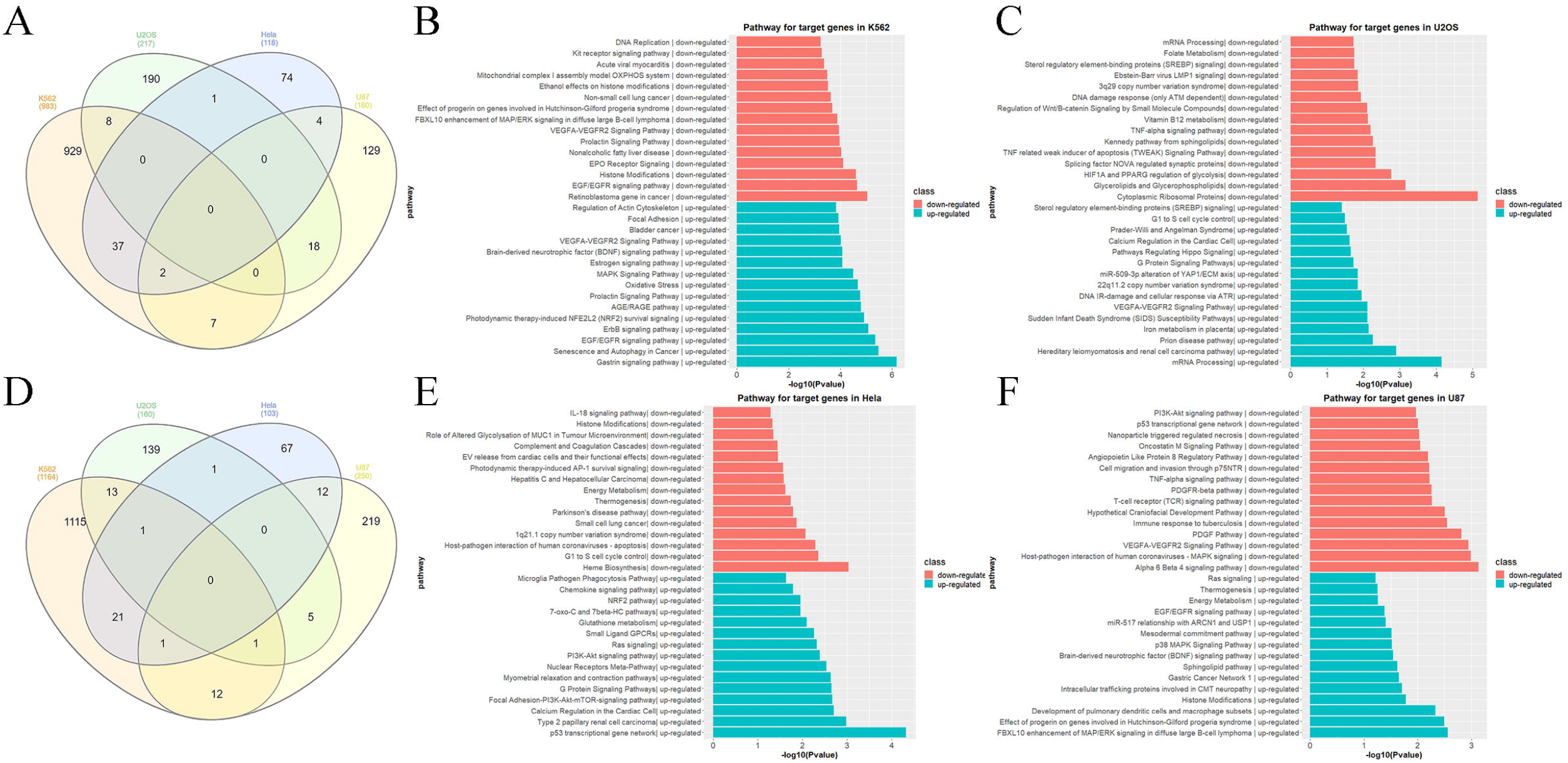
Transcription factors that are down-regulated by R-loops across cell lines.

